# Relentless Selection: The importance of within-generation selection in heterogeneous habitats

**DOI:** 10.1101/2023.12.26.573350

**Authors:** Moritz A. Ehrlich, Amanda N. DeLiberto, Melissa K. Drown, Marjorie F. Oleksiak, Douglas L. Crawford

## Abstract

Natural selection relentlessly reshapes the genetic and phenotypic composition of populations, yet often adaptations cannot emerge due to excessive migration and gene flow. Nevertheless, in heterogeneous habitats strong selection could temporarily establish significant trait divergence among environmental patches. Here, we show that in *Fundulus heteroclitus,* a single generation of selection drives significant phenotypic divergence (5-15%) in organismal metabolic rate, cardiac metabolic rate and hypoxia tolerance. This divergence occurs among individuals of the same panmictic population residing in environmentally distinct microhabitats. Phenotypic divergence remains observable following long-term common-gardening and is supported by previous work documenting fine-scale, genetic divergence among microhabitat residents. We show that the magnitude of within-generation trait divergence is on the order of what is commonly observed among more isolated populations that have diverged over multiple generations. Although panmictic reproduction among microhabitat residents erodes trait divergence every generation, strong selection could potentially reestablish it in the next. In heterogeneous habitats, transient, fine-scale divergence could have a considerable impact on eco-evolutionary dynamics. Ignoring its contribution to overall trait variance could limit our ability to define meaningful, evolved divergence.

**Summary:** Natural selection can lead to changes in organisms’ traits over time. Typically, these changes occur slowly over multiple generations and over large spatial scales. By studying a wild population of Atlantic killifish, we show that a single generation of natural selection can generate substantial trait variation over short distances. We observe significant differences in several physiological traits among individuals inhabiting distinct ‘microhabitats’ in a patchy salt marsh environment. These differences are unlikely due to physiological acclimation and are best explained by strong, natural selection removing those individuals not suited to a particular microhabitat. Previous studies support natural selection as the most likely explanation, having shown subtle genetic differences among microhabitat residents. Remarkably, the magnitude of trait divergence is on the order of what is typically observed among populations that have diverged over multiple generations and larger spatial scales. Our results highlight the significant contribution of natural selection to trait variation in patchy environments, even over exceptionally short time and small spatial scales.

## Introduction

Comprehending the importance of natural selection in driving evolution is critical to our understanding of natural systems and how these will be affected by future changes to the environment (Hendry and Gonzalez 2008). Natural selection can result in long-term evolutionary change: when its effects compound over several generations, continued selection can lead to significant phenotypic alterations that increase the relative fitness of a population. While adaptive evolution was historically viewed as a slow process, requiring on the order of *N_e_* generations even under strong selection (Otto and Whitlock 2013), recent findings suggest it may be more effective than previously thought. Organisms seem to be evolving and adapting on much shorter time scales and smaller spatial scales than classic evolutionary theory predicts (Richardson et al. 2014; Messer et al. 2016; Reznick et al. 2019; Grainger and Levine 2022). Despite its apparent effectiveness, adaptation on small spatial scales is often inhibited by excessive gene flow (Lenormand et al. 2002; Yeaman and Whitlock 2011) or strong drift when selection is weak (*s* < 1/2*N*_e_)(Kimura 1962). Under high migration rates or unfavorable admixture, adaptive trait divergence generated by selection cannot propagate into the next generation, resulting in net-zero change and effectively inhibiting adaptative evolution (Felsenstein 1976). Nevertheless, there is evidence to suggest selection may be effective enough to repeatedly reestablish significant trait divergence every generation. For instance, individuals belonging to a genetically unstructured population may reside in distinct microhabitats and experience opposing selection pressures over their lifetime. Natural selection may then lead to temporary phenotypic divergence among microhabitats, subsequently erased by panmictic reproduction (Gagnaire et al. 2012; Laporte et al. 2016; Rey et al. 2020). Although transient, this momentary divergence can result in higher trait variance than expected in unstructured populations and could potentially be more ecologically relevant than global acclimation responses or long-term adaptive changes. Here, we quantify transient, within-generation divergence of ecologically important physiological traits among residents of distinct microhabitats. By sampling a natural, unstructured and interbreeding population of Atlantic killifish (*Fundulus heteroclitus*), we demonstrate that a single generation of selection generates significant trait divergence among residents of distinct microhabitats.

*F. heteroclitus* is a small killifish inhabiting saltmarsh estuaries along the eastern coast of the USA. It is a keystone species consisting of populations typically exceeding 10,000 individuals (Duvernell et al. 2008) and reaching biomass densities of 160 kg/ha dry weight (Valiela et al. 1977). Due to its high productivity, it is also a major food source for a number of aquatic and avian predators (Kneib 1986). *F. heteroclitus* has been shown to exhibit remarkable tolerance to changes in temperature, salinity, dissolved oxygen concentration and even toxins, allowing it to inhabit a wide range of habitats (Crawford et al. 2020). Within saltmarsh estuaries *F. heteroclitus* inhabit several microhabitats such as shallow marsh ponds, tidal channels and deeper coastal basins. Notably, environmental differences among microhabitats within the same estuary can be particularly stark. During the summer months shallow ponds with a mean surface area of 100 m^2^ and a mean depth of only 40 cm can occasionally reach temperatures above 38 °C. In addition, high photosynthetic activity can hyper-oxygenate ponds during daytime (>20 mg/L) whilst net respiration at night frequently leads to complete anoxia for several hours (Smith and Able 2003; Hunter et al. 2009). Heavy rains and wind-driven flooding can also cause sudden shifts in pond temperature, salinity and oxygen concentration (Sidell et al. 1983). These challenging conditions, especially during the summer months, lead to mortality in excess of 30% per month for *F. heteroclitus* inhabiting marsh ponds (Hunter et al. 2009). In contrast to the extreme and highly variable pond environment, coastal basins present a more stable microhabitat. These deeper coastal inlets continually exchange water with the open sea, maintaining relatively stable temperature, salinity and oxygen profiles. Nevertheless, coastal basins likely exhibit other selective pressures such as higher aquatic predation and competition for food (Kneib 1986).

Although *Fundulus heteroclitus* inhabit a highly connected landscape, individual fish display remarkable site fidelity (Lotrich 1975; Skinner et al. 2005; Able et al. 2012). Despite strong tidal flows, individuals display a limited home range, even returning to their respective home ranges after being released at a different location (Lotrich 1975). Using implant tags, Skinner et al. (2005) showed more than 95% of *F. heteroclitus* remain within 200 m of their tagging location after one year. In fact, over 90% of individuals appear to reside within distinct microhabitats for several months (Wagner et al. 2017; Ehrlich et al. 2021). Given high site fidelity and minimal migration one might expect substantial demographic structure, yet *F. heteroclitus’* reproductive cycle ensures strong admixture among all individuals inhabiting the same estuary (Taylor et al. 1979). During spring tides (highest tides in the lunar cycle) salt marshes flood entirely, and individuals from all microhabitats congregate at the high-water line to mate (Taylor et al. 1979). This synchronized reproduction at common breeding grounds leads to *F. heteroclitus* forming a single, panmictic population with large effective population size on the order of 10^4^ individuals (Duvernell et al. 2008) and with no genetic structure among microhabitats (Ehrlich et al. 2021).

We harness the *F. heteroclitus* system in which individuals belonging to the same panmictic population inhabit highly contrasting natural environments. We show that selection is strong enough to generate significant trait divergence between pond and basin residents over the course of just a single generation. This divergence occurs in complex physiological traits that have been associated with adaptation to different temperature and oxygen regimes (Oleksiak et al. 2005; Schulte 2015; Pörtner et al. 2017; Dayan et al. 2019; Crawford et al. 2020). This strong, within-generation selection produces fine-scale trait heterogeneity across microhabitats that is unexpected given panmictic breeding. Importantly, we show that within-generation selection generates significant trait divergence on the order of that caused by long-term evolution among more isolated populations.

## Results

### High site fidelity

After tagging over 2,300 *F. heteroclitus* individuals in May 2018 in three marsh ponds and one coastal basin, we successfully recaptured a total of 271 individuals across the four sampling sites in September 2018 (Fig. 1). Sampling effort for recapture was extensive, trapping nearly all individuals in the target ponds and sampling the coastal basin until less than one tagged fish was trapped per 15 wire trap deployments. Recapture rates ranged from 9.5% to 14.5% with 131 (Basin), 58 (Pond 1), 44 (Pond 2) and 38 (Pond 3) fish recaptured at each site. Tag data confirmed high site fidelity with 89% of individuals recaptured at their respective tagging location. Residency rates were comparable among sites (Basin 86%; Pond 1 89%; Pond 2 88%; Pond 3 100%). Physiological measurements were only analyzed for resident individuals that had been exposed to their respective microhabitat’s ambient conditions from spring to late summer.

**Fig 1.**
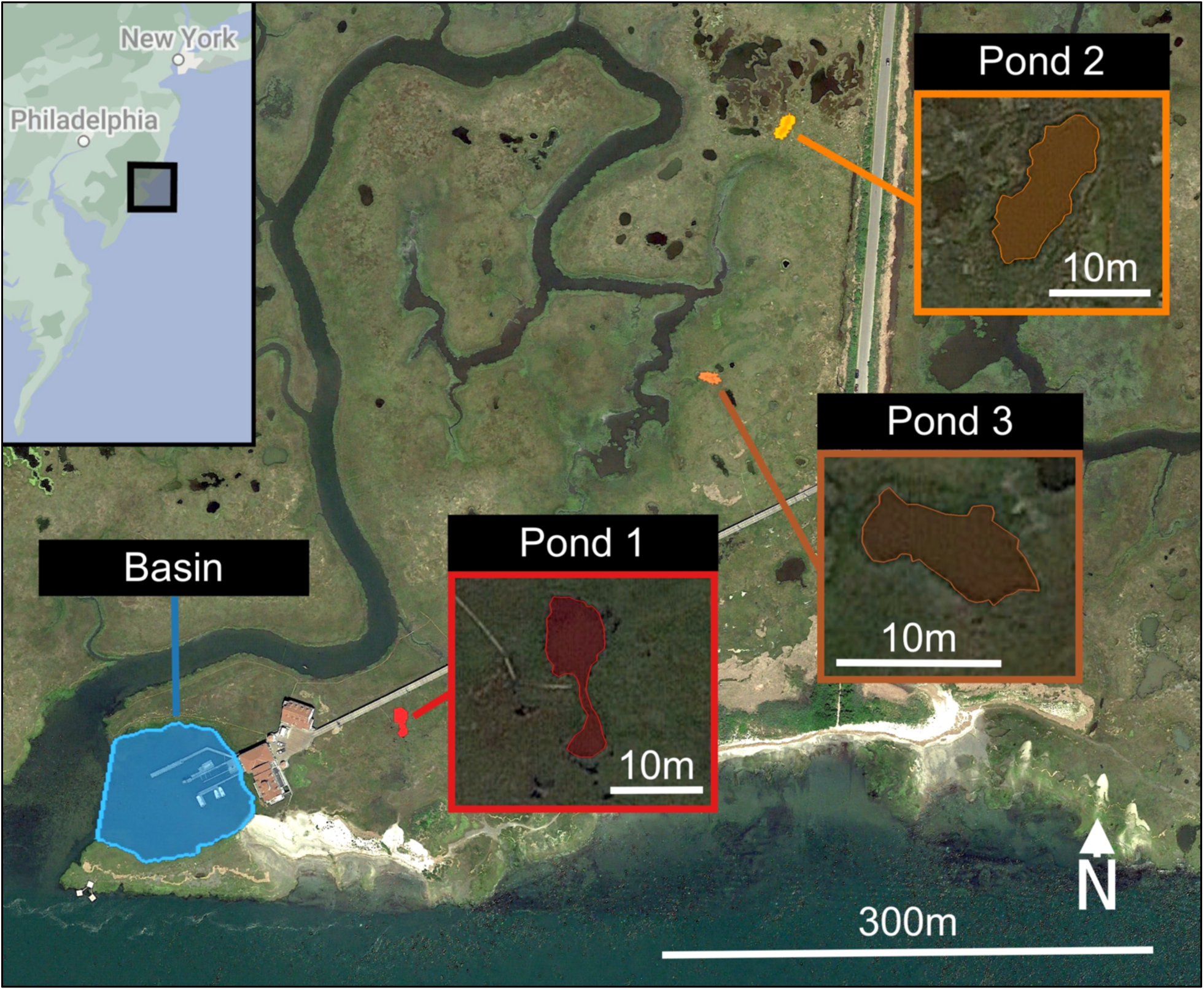
Sampling Locations. Satellite image of the four sampling locations near the Rutgers University Marine Field Station in Little Egg Harbor, New Jersey. Environmental data was collected, and fish were extracted from one coastal basin and three marsh ponds within the same estuary.

### Distinct microhabitats

Continuous deployment of environmental data loggers during summer 2018 confirmed highly disparate conditions between the coastal basin and the marsh ponds (Fig. 2)(Smith and Able 2003; Hunter et al. 2009). Marsh ponds showed significantly higher daily mean temperatures than the coastal basin (+3.9 °C, s.d. = 2.0)(Fig. 2A and Supplement 1). Since ponds are smaller, isolated bodies of water, we also observed greater diurnal variation in temperature with higher frequency and duration of temperature peaks compared to the basin. Specifically, we recorded temperature extremes above 28°C on 63% of days in every pond, some lasting over 12 hours (Fig. 2B). In contrast, coastal basin temperatures exceeded 28°C on only 10% of measured days. When temperatures did exceed 28°C in the basin, the duration was short, typically lasting 5 hours or less (Supplement 1).

**Fig 2.**
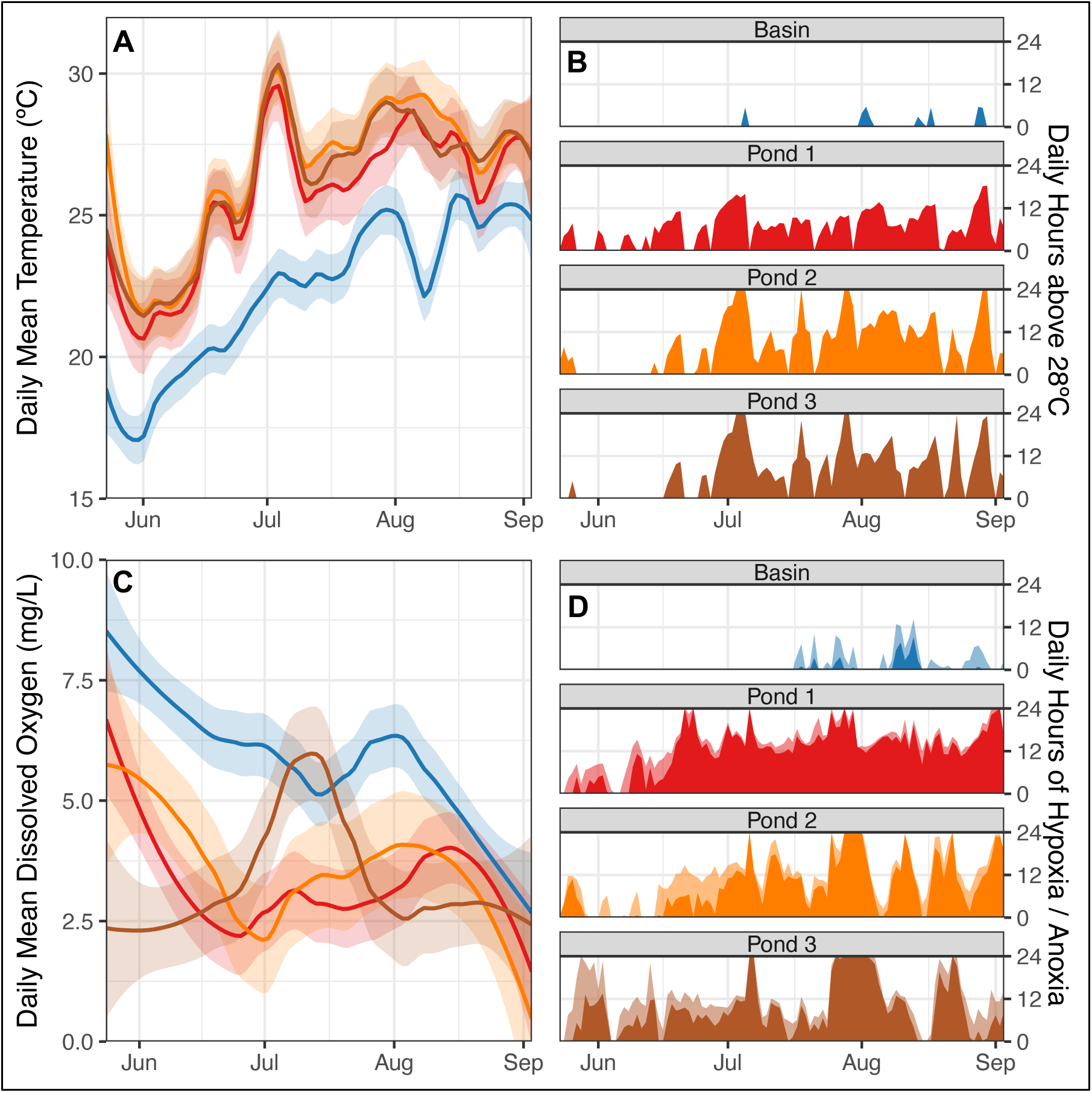
Environmentally distinct microhabitats. (A) Smoothed daily mean temperature of the coastal basin (blue) and the three marsh ponds (shades of red). Shaded bands show the standard deviation about the mean. (B) Total daily hours during which temperatures exceeded 28°C at each location. (C) Smoothed daily mean dissolved oxygen concentration of the coastal basin (blue) and the three marsh ponds (shades of red). Shaded bands show the standard deviation about the mean. (D) Total daily hours during which hypoxic (< 2 mg/L, light shading) and anoxic (< 0.5 mg/L, dark shading) conditions were recorded at each location.

Even more striking than the disparate temperature regimes are the differences in dissolved oxygen concentration: marsh ponds exhibit significantly lower daily mean dissolved oxygen relative to the coastal basin (-2.3 mg/L, s.d. = 2.3, Fig. 2C). Ponds also display larger diurnal variation in oxygen concentration. Prolific photosynthesis during the day leads to supersaturation with oxygen concentrations exceeding 20 mg/L. At night net respiration rapidly depletes oxygen, leading to prolonged hypoxic periods (< 2 mg/L) and even anoxia (< 0.5 mg/L) in the early hours of the morning. While these hypoxic periods typically lasted for 12 hours, our observations show they can occasionally extend over several days in the marsh ponds (Fig. 2D). The coastal basin on the other hand did not experience these extreme fluctuations. Oxygen concentrations were moderately high throughout our measurement period (5.76 mg/L, s.d. = 2.83), and hypoxic conditions were rare and of short duration (Fig. 2D).

### Trait Divergence

Recaptured pond and basin residents were common-gardened in a closed, recirculating system for over 8 months to minimize prior acclimation effects. After common-gardening, seven physiological traits were quantified: 1) organismal standard metabolic rate (SMR), 2) aquatic surface respiration (ASR) as a proxy for hypoxia tolerance, 3) critical thermal maximum (CT_max_) as a proxy for thermal tolerance and 4-7) cardiac metabolic rate under four substrate conditions: i) glucose, ii) fatty acids, iii) lactate, ketones and alcohol (LKA) and iv) no substrate (endogenous metabolic rate). We observed significant divergence in three of these complex traits: organismal SMR, cardiac metabolic rate with glucose substrate, and ASR were significantly different between basin and pond residents (Fig. 3).

**Fig 3.**
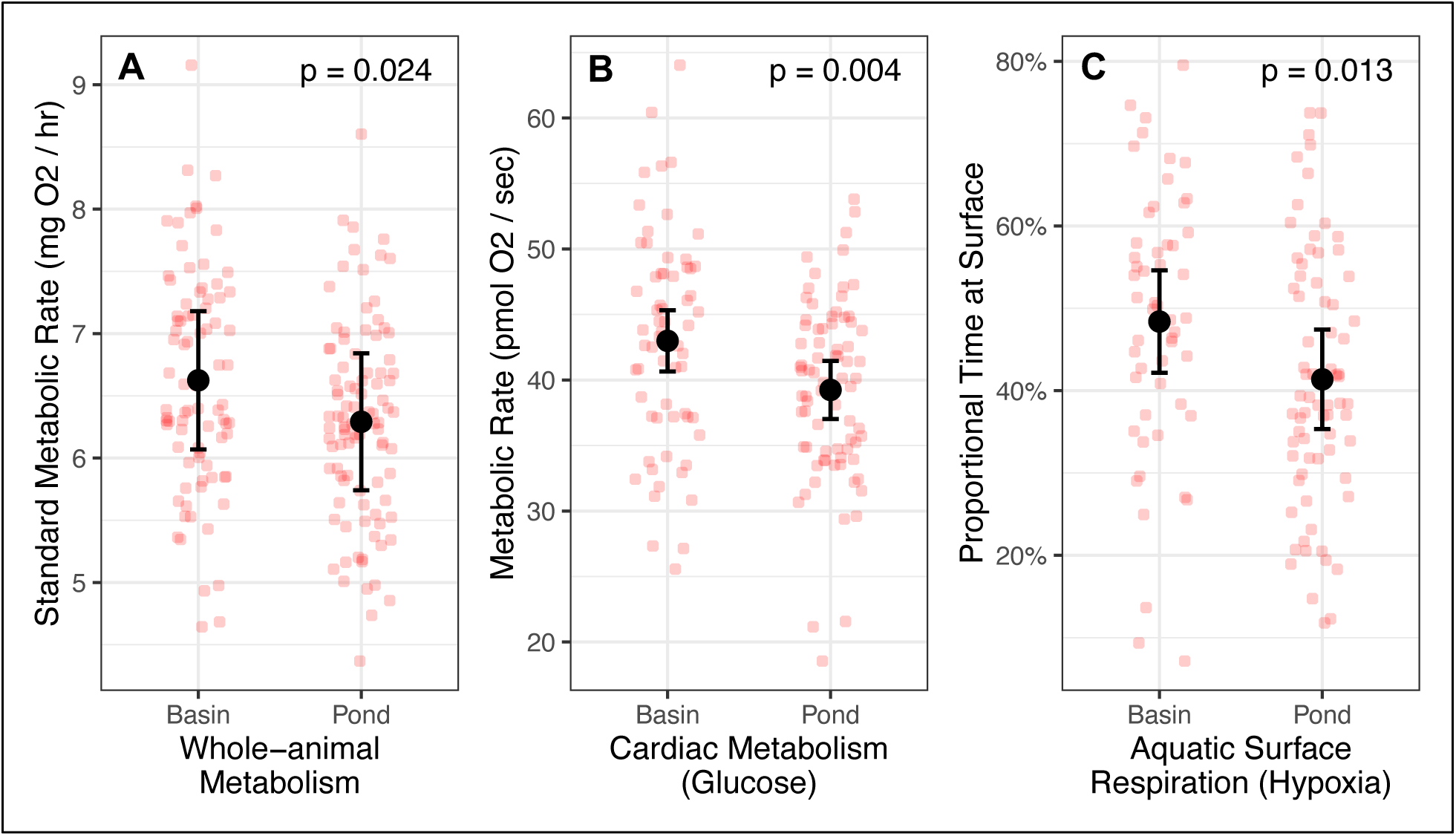
Significant physiological differences among residents of distinct microhabitats. Significant divergence in three physiological traits between coastal basin and marsh pond residents measured at 28°C. Black circles show the marginal means of the respective linear mixed model, error bars show the standard error about the mean, and red circles show individual data points adjusted for covariates. (A) Significant difference in whole-organism standard metabolic rate between basin and pond residents. (B) Significant difference in cardiac metabolic rate between basin and pond residents with glucose as the principal substrate. (C) Significant difference in the duration of aquatic surface respiration during hypoxia between basin and pond residents. Black circles show the marginal means of the respective linear mixed model, error bars show the standard error about the mean, and red circles show individual data points adjusted for covariates.

Pond residents exhibited a 5.0% lower SMR than basin residents after correcting for body mass (*p* = 0.024, Fig. 3A). As expected, organismal mass was a highly significant covariate (*p* << 0.05) of SMR while none of the other potential covariates (e.g. sex) nor their interactions were significant (Supplement 2). Lower SMR in ponds as opposed to basins is congruent with the theory of oxygen and capacity-limited thermal tolerance (OCLTT). It proposes that at high temperatures performance is constrained by the inability to deliver sufficient oxygen to relevant tissues (Pörtner et al. 2017). In marsh ponds this is further exacerbated by low ambient oxygen concentrations. As a result, individuals with a relatively high SMR may be selected against in hot, oxygen-limited ponds where higher basal metabolic demands restrict the aerobic scope for e.g. foraging or predator evasion (Hochachka and Somero 2014). In contrast, individuals with a lower SMR may be selected against in the basin where oxygen is more abundant and lower basal metabolism is associated with slower growth rates and decreased intraspecific competitiveness (Metcalfe et al. 2016).

Cardiac metabolic rate with glucose as a substrate showed a similar pattern to SMR. Pond residents displayed 8.7% lower cardiac metabolic rate compared to their basin counterparts (*p* = 0.004, Fig. 3B). Again, this may be the result of selection against individuals with higher metabolic demands in hot, hypoxic pond environments, and against individuals with lower metabolic rates in the more densely populated basin where intraspecific competition could be higher. Cardiac mass was a significant covariate (*p* << 0.05) as was assay time due to oxygen sensor drift during measurement sessions. However, the systematic effect of sensor drift was accounted for in the linear mixed model, and assay order of basin and pond individuals was randomized to further mitigate sensor drift effects.

Pond residents spent 14.5% less time performing ASR under hypoxic conditions (< 2mg/L) than basin residents (*p* = 0.013, Fig. 3C). Using ASR duration as a proxy for hypoxia tolerance, our results suggest pond residents are more tolerant of hypoxic conditions. The linear mixed model also showed organismal mass and sex to be significant covariates of ASR duration with smaller, female fish spending significantly more time performing ASR than larger males. A separate multi-state model analysis, which models the instantaneous probability of an individual transitioning between the submerged and ASR state as a function of dissolved oxygen, also supports these results (Fig. 4). Pond individuals are significantly less likely to initiate ASR at intermediate or low oxygen concentrations than their basin counterparts (solid lines, Fig. 4). In addition, pond residents are also significantly more likely to terminate ASR than basin residents (dashed lines, Fig. 4). In consequence, single ASR events are both less frequent and shorter in pond residents suggesting higher hypoxia tolerance.

**Fig 4.**
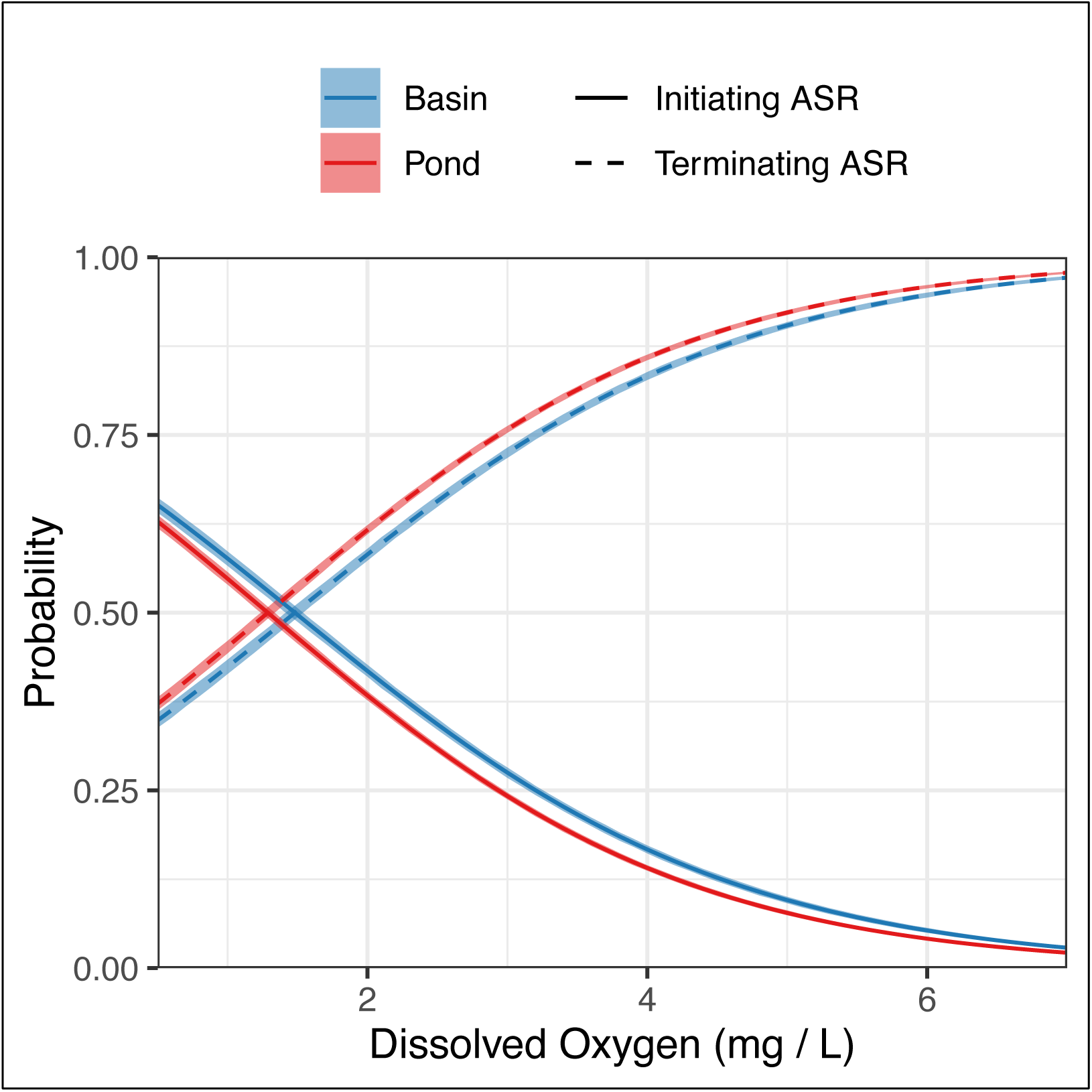
Significant difference in ASR behavior among basin and pond residents. Transition probabilities between the ASR and non-ASR state extracted from the multi-state model. Colored lines show the instantaneous probability of an individual to initiate ASR if submerged (solid line) or terminate ASR if surfaced (dotted line) as a function of dissolved oxygen. Shaded bands show the 95% confidence interval. Pond individuals (red) are significantly less likely to initiate ASR and significantly more likely to terminate ASR than basin individuals (blue).

Whilst we observed significant differences between pond and basin residents in three complex physiological traits, other measured traits did not show significant divergence. Residents of the two microhabitats did not differ in their CT_max_ (*p* = 0.14, Supplement 2) with none of the covariates showing a significant effect. Cardiac metabolic rate with a fatty acid substrate showed marginal divergence (*p* = 0.053). Again, pond residents exhibited a lower cardiac metabolic rate than basin individuals, congruent with organismal SMR and glucose cardiac metabolism. Differences in LKA cardiac metabolic rate (*p* = 0.437) and endogenous cardiac metabolic rate (*p* = 0.172), were insignificant, but trended in the same direction with pond residents displaying lower metabolic rates (Supplement 2).

Substrate-specific cardiac metabolic rates were significantly correlated (*p* << 0.05) with moderate Pearson correlation coefficients (0.33 ≤ *r* ≤ 0.47). None of the other physiological traits were significantly correlated except for a weak association between ASR latency and endogenous cardiac metabolic rate (*r* ≤ 0.26, *p* = 0.01)(Supplement 3).

## Discussion

### Within-generation selection causes significant trait divergence

We propose that the significant trait differences between residents of distinct microhabitats are the result of natural selection acting over a single generation. These metabolic trait differences are observable after 8 months of common-gardening, suggesting that they have a genetic basis rather than being driven by physiological plasticity. This is further supported by previous work showing significant allele frequency changes occurring in both basin and pond residents during summer when mortality is highest (Hunter et al. 2009; Ehrlich et al. 2021).

Allele frequency changes were unlikely due to random mortality or migration and resulted in fine genetic structure at specific loci, segregating basin and pond residents (Wagner et al. 2017; Ehrlich et al. 2021). Moreover, data collected from previous generations of *F. heteroclitus* in the same estuary suggest that natural selection could potentially generate trait divergence repeatedly every generation. High site fidelity in *F. heteroclitus* exposes individuals of the same interbreeding population to distinct microhabitats with unique phenotypic optima. Distinct selection pressures consequently drive strong selection that generates significant divergence in SMR, cardiac metabolic rate and hypoxia tolerance. This results in a heterogeneous trait distribution across habitat space where trait variation is associated with microhabitat. However, this trait divergence is most likely transient as synchronous reproduction at common breeding grounds homogenizes allele frequencies leading to complete admixture across the estuary (Ehrlich et al. 2021). Genetically homogeneous offspring distribute themselves among microhabitats and the process of within-generation selection repeats.

While repeated, within-generation selection could lead to the observed trait differences between basin and pond residents, there might be alternative explanations. Developmental effects, whereby the local environment experienced by juveniles during maturation irreversibly conditions their phenotype, could also explain our results. Differential exposure to certain temperatures, salinities, or compounds at the larval stage could lead to alternative development or epigenetic modifications, that can potentially cause physiological differences in adult fish, even after common-gardening (Jonsson and Jonsson 2014). In fact, the effect of prior exposure on current phenotype is not limited to the juvenile stage. The environment experienced by parental fish could also affect offspring phenotype via epigenetic inheritance (Anastasiadi et al. 2021). However, since neither alternative development nor epigenetic changes modify the underlying genotype, prior exposure cannot account for the significant allele frequency changes previously detected during adulthood (Ehrlich et al. 2021). In addition, stable epigenetic inheritance would require assortative mating between residents of the same microhabitat. This would generate significant genetic structure which has not been observed (Wagner et al. 2017; Ehrlich et al. 2021).

Another potential explanation is that pond residents display higher hypoxia tolerance because of learned behavior. Repeated exposure to hypoxia in the wild may have allowed pond residents to optimize their technique to allow for shorter and less frequent ASR bouts. Similar to developmental effects, learned behavior cannot be controlled by common-gardening. Still, learned behavior is unlikely to generate differences in metabolic rates, especially at the organ level as was the case for cardiac metabolism.

Finally, it is possible that the observed trait differences are due to the deliberate movement of individuals seeking out favorable environmental conditions (Edelaar et al. 2008; Nicolaus and Edelaar 2018). Matching habitat choice, the concept of individuals actively sorting themselves into microhabitats that maximize their fitness relative to their genotype, has been observed in ciliates (Jacob et al. 2017), grasshoppers (Camacho et al. 2020), salamanders (Lowe and Addis 2019), and three-spine stickleback (Bolnick et al. 2009). This phenomenon could explain both trait divergence as well as the allele frequency shifts in pond and basin residents. *F. heteroclitus*, however, is severely limited in its ability to migrate between microhabitats since infrequent flooding of the marsh only gives temporary access to the ponds (Hunter et al. 2009). In addition, the marsh environment is homogenized during flooding periods such that environment-dependent migration is unlikely to occur. This is further supported by previous studies that have reported limited migration patterns in *F. heteroclitus,* indicating high site fidelity and restricted home ranges (Lotrich 1975; Skinner et al. 2005). Considering i) panmictic reproduction within the estuary (Taylor et al. 1979) ii) high site fidelity (Lotrich 1975; Skinner et al. 2005) iii) high mid-summer mortality (Hunter et al. 2009) and iv) non-random allele frequency changes that generate subtle genetic divergence (Wagner et al. 2017; Ehrlich et al. 2021); within-generation natural selection seems to be the most parsimonious explanation for the observed trait divergence.

### The importance of within-generation selection

Even strong selection is unlikely to lead to long-term, local adaptation if reproduction is panmictic among microhabitats. For locally adapted phenotypes to evolve, divergence generated by natural selection must propagate to the next generation and accumulate over time (Hendry and Gonzalez 2008). In *F. heteroclitus* phenotypic divergence between residents of distinct microhabitats is likely to recede each generation because individuals form a single, panmictic population. This characterization is supported by synchronized reproduction at common breeding grounds (Taylor et al. 1979) and by the lack of population structure (Wagner et al. 2017; Ehrlich et al. 2021). Repeated and complete swamping by maladapted alleles from a panmictic reproductive pool limits local adaptation. Migration-selection balance disfavors long-term population divergence and local adaptation is not expected to evolve (Lenormand et al. 2002; Yeaman and Whitlock 2011; Tigano and Friesen 2016). Thus, while strong selection may generate significant phenotypic divergence in a single generation, any divergence between basin and pond residents is likely reset in the next generation. Long-term evolutionary changes are only expected to occur at the global level i.e. across all microhabitats, and in response to the geometric mean fitness integrated over time (Connallon and Clark 2012; Yeaman 2015; Bertram and Masel 2019). For example, widespread temperature increases due to climate change could drive a global adaptive response across all microhabitats.

While locally adapted phenotypes are unlikely to evolve under these conditions, the effect of within-generation selection should not be dismissed only because it is temporary. Figure 5 shows the standardized trait divergence between pond and basin residents resulting from a single generation of selection (this study – red triangles) and compares it to the distribution of standardized trait divergence resulting from long-term evolution, observed between natural populations as published in the literature (Supplement 4 and 5). Data were collected from publications documenting significant, heritable trait divergence between two or more natural populations of the same species (Supplement 4 and 5). Remarkably, trait divergence caused by selection over a single generation in *F. heteroclitus* is comparable in magnitude to long-term evolutionary divergence across much greater spatial and temporal scales. Although our results fall into the lower quadrant of the distribution, within-generation divergence in hypoxia tolerance and cardiac metabolic rate is larger than 15% of evolved trait differences among other natural populations. Moreover, this number is likely to be a conservative estimate, given the ascertainment bias in documenting small yet significant effect sizes (Hendry 2013).

**Fig 5.**
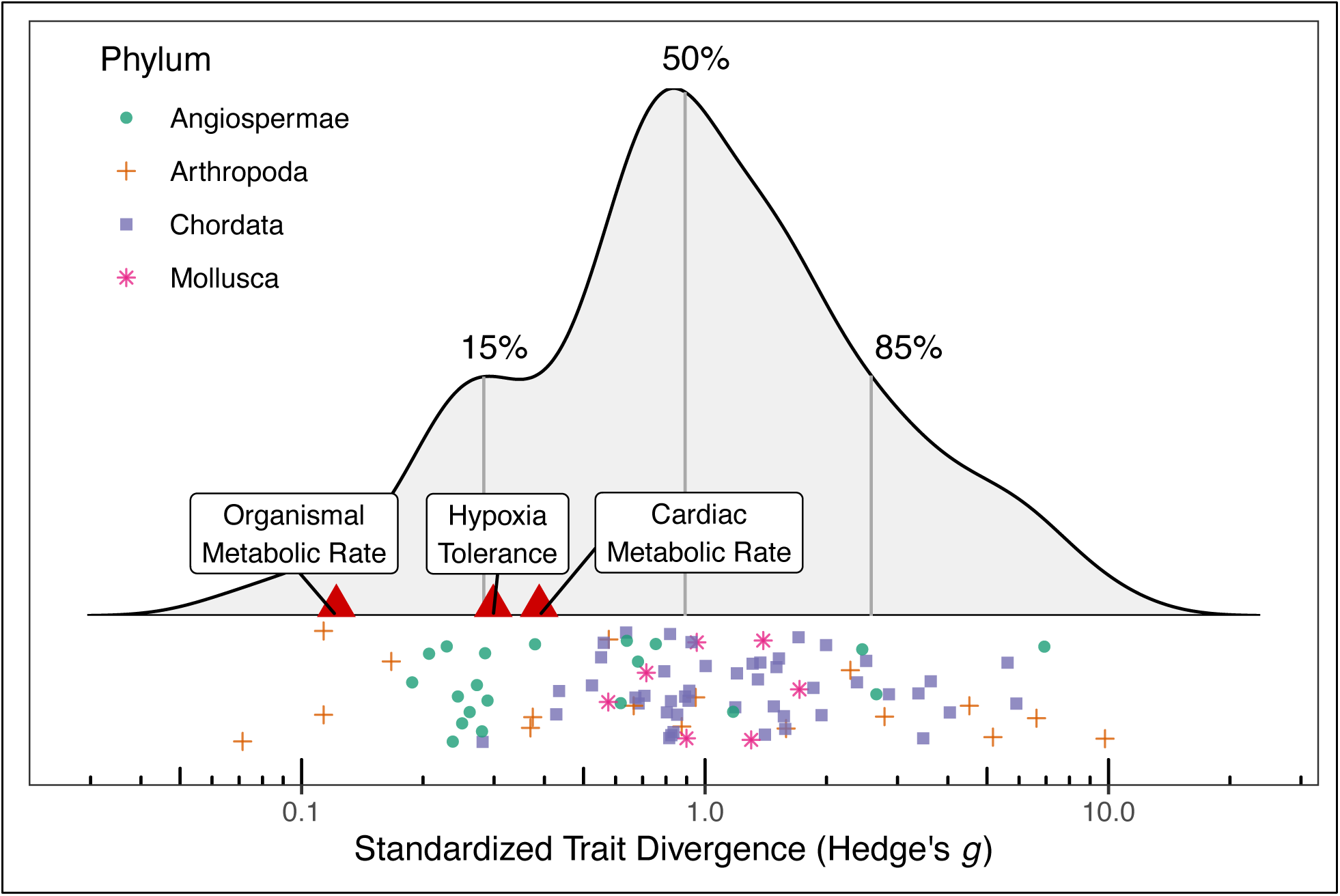
Within-generation selection generate significant trait divergence comparable to that of local adaptation. Distribution of standardized trait divergence between pairs of locally adapted populations, colored by phylum. Data were retrieved from publications documenting significant, heritable divergence among natural populations by means of common gardening or reciprocal transplant designs (Supplement 4 and 5). Each data point represents the standardized trait divergence of a given trait between a given population pair. Vertical lines show the 15^th^, 50^th^, and 85^th^ percentiles. Red triangles show the standardized trait divergence between pond and basin residents resulting from a single generation of selection.

Our results highlight how a single generation of strong selection could generate significant spatial trait variation within an interbreeding population. In heterogeneous habitats, this transient divergence could potentially be of considerable importance on shorter, ecological time scales (Hendry 2013; Urban et al. 2020). For example, several species of wading bird typically prey on *F. heteroclitus* in hypoxic marsh ponds as fish are forced to the surface. Wading birds account for a significant proportion of *F. heteroclitus* mortality (Kneib 1986) and could be important selective agents. By preferentially removing individuals that surface more frequently, wading birds could be responsible for the divergence in hypoxia tolerance between basin and pond residents. In turn, wading bird foraging success, and therefore their distribution, could be impacted by the spatial distribution of hypoxia tolerance in *F. heteroclitus*. That is, birds are more likely to forage where they have a higher success rate, which is where fish have the lowest oxygen tolerance relative to ambient conditions. Hence, within-generation selection and the resulting spatial heterogeneity could potentially impact key ecosystem features like predator distribution.

Ignoring the contribution of within-generation selection can also lead to bias by underestimating the trait variation among microhabitats. This unaccounted variance within a population can potentially confound estimates of between-population variance and reduce our ability to define significant, evolved divergence. For example, sampling a different microhabitat in each of two isolated populations could result in inflated estimates of population divergence and biased conclusions regarding the extent of local adaptation. This bias is likely to be most pronounced in panmictic populations inhabiting heterogeneous landscapes (Richardson et al. 2014; Bernatchez 2016; Denney et al. 2020) such as sessile, broadcast-spawning organisms like cross-pollinating plants (Antonovics 2006) or certain marine invertebrates (Nunez et al. 2021), and mobile organisms with common breeding grounds yet high site-fidelity to disparate niche environments (Gagnaire et al. 2012).

### Maintaining genetic variance

Strong selection occurring repeatedly every generation poses a non-trivial problem. Natural selection requires genetic variance to act upon, yet persistent, strong selection is expected to significantly reduce genetic variance over time. Without a mechanism to generate or maintain genetic variance, repeated within-generation selection is unsustainable. Given the extremely short time scale of within-generation selection, new mutations are unlikely to be a meaningful source of genetic variance. Similarly, evidence showing low migration rates between disconnected estuaries suggests immigration contributes little to genetic variation (Duvernell et al. 2008).

Balancing selection offers a potential solution to the challenge of maintaining genetic variance. Analogous to Levene’s (1953) two-patch model with complete admixture, alleles showing approximately equal and opposite selection coefficients in the two microhabitats could theoretically be maintained at intermediate frequencies indefinitely (Gillespie 1974). Opposing selection coefficients may arise for traits that are beneficial in one and deleterious in the other environment or through antagonistic pleiotropy of two or more traits (Brown and Kelly 2018). For instance, a lower standard metabolic rate may be beneficial in hypoxic pond environments due to lower oxygen demand (Hochachka and Somero 2014). In coastal basins, however, higher SMR may allow for faster growth or improve intraspecific competition (Metcalfe et al. 2016). In the case of antagonistic pleiotropy different traits with an overlapping genetic basis may be under selection in the two environments. This generates genetic conflict that can potentially maintain allele frequencies at intermediate values (Brown and Kelly 2018).

Regardless of the mode of balancing selection, the conditions required for the stable maintenance of polymorphism in heterogeneous environments are extremely limiting (Hedrick 1999; Akerman and Bürger 2014). Natural populations are unlikely to experience these restrictive conditions and few examples of balancing selection have been documented so far (but see Gagnaire et al., 2012; Bergland et al., 2014; Nunez et al., 2021). In most cases, a generalist genotype exhibiting the highest geometric mean fitness over time and across microhabitats is expected to establish, and this will erode genetic variation (Haldane and Jayakar 1963).

Despite these limitations, recent work has shown that, depending on the genetic architecture, conditions for the maintenance of polymorphism may not be as restrictive as previously thought, especially when considering shorter time scales (Connallon and Clark 2012; Yeaman 2015). The compound, physiological traits measured here, were previously shown to have complex, polygenic architectures with high redundancy and small per-locus effect sizes (Shi et al. 2016; Barghi et al. 2019). Selection on polygenic traits can lead to significant trait shifts, yet inherent redundancy among the underlying loci implies that the probability of any one allele going to fixation (“hard sweep”) is low (Höllinger et al. 2019). In finite populations, alleles underlying polygenic traits tend to fix, and hence be removed, more slowly (Thornton 2019) with the time to fixation (and loss) of an allele on the order of *N_e_* (Kimura and Ohta 1969; Thornton 2019). Genetic variance is therefore eroded more slowly in large populations under polygenic selection (Lande 1975; Höllinger et al. 2019).

Loss of polymorphism can be slowed further in the presence of antagonistic pleiotropy. Antagonistic alleles are expected to be fixed (or removed) more slowly than unequivocally beneficial (or deleterious) alleles (Lynch and Walsh 1998). Connallon & Clark (2012) showed that under antagonism the rate of allele frequency change is proportional to the product of environment-specific selection coefficients. Assuming small selection coefficients, the rate of fixation, and thus the rate of loss of polymorphism, can be several orders of magnitude lower than for universally beneficial or deleterious alleles (Connallon and Clark 2012). In summary, polygenic architectures and antagonistic pleiotropy could slow the loss of polymorphism under selection and allow genetic variance to be maintained over longer periods.

It should be emphasized that these processes are not stable equilibria, and that the indefinite maintenance of polymorphism remains unlikely. The evolution of a generalist genotype, modifier genes, or reproductive isolation between basin and pond residents are more probable outcomes in the long-term (Yeaman and Whitlock 2011; Yeaman 2015). Rather, polygenic architectures with high redundancy should be viewed as extending the time until such equilibria are reached. Yeaman (2015) showed that for polygenic selection with high migration, significant genetic variance can be maintained for time periods on the order of 10^4^ generations. In the case of *F. heteroclitus*, the end of the last glaciation event dates back approximately 10^4^ generations. Following glacial retreat, contact between previously isolated populations (Haney et al. 2009) might have generated substantial standing genetic variation in *F. heteroclitus*. High standing variation, large population sizes and polygenic architectures could potentially be fueling repeated, within-generation selection and the resulting, transient divergence between pond and basin residents.

## Conclusion

We have quantified significant divergence (5-15%) in three complex, physiological traits among *F. heteroclitus* individuals that belong to the same unstructured population yet reside in environmentally distinct microhabitats. This trait divergence is unlikely an acclimatory response but most likely the result of a single generation of selection. Previous studies support this interpretation, showing subtle genetic divergence among residents of distinct microhabitats (Wagner et al. 2017) caused by significant, within-generation allele frequency changes over the summer season when mortality is highest (Ehrlich et al. 2021). Panmictic reproduction at common breeding grounds homogenizes geno- and phenotypes, suggesting trait divergence among microhabitat residents is temporary. Nevertheless, the magnitude of within-generation divergence is on the order of what is commonly observed among more isolated populations that have evolved over longer timeframes. In heterogeneous habitats, this transient, fine-scale divergence could potentially be of considerable importance on shorter, ecological time scales (Hendry 2013; Urban et al. 2020). Particularly with global climate change contributing to increased environmental heterogeneity, the impact of within-generation selection could intensify. Ignoring the contribution of within-generation selection to trait variance could lead to substantial bias by underestimating variation among microhabitats. Unaccounted within-population variance can potentially confound estimates of between-population variance, reducing our ability to define significant, evolved divergence. Lastly, repeated, strong selection requires sufficient standing genetic variation to act upon. Large population sizes, polygenic architectures, and antagonistic pleiotropy could potentially slow the rate of loss of polymorphism making repeated, within-generation selection feasible in short to medium evolutionary terms.

Although previous work supports within-generation, divergent selection (Wagner et al. 2017; Ehrlich et al. 2021), we acknowledge that our suggestions regarding repeated trait divergence between pond and basin *F. heteroclitus* as well as our potential explanation for the maintenance of genetic variance are speculative and require further testing. Whole-genome sequencing data at within-generation resolution could uncover the genetic architecture of within-generation divergence through measurements of allele frequency changes in pond and basin residents. Furthermore, cross-referencing alleles responding to selection with those associated with the physiological traits measured here could yield target loci involved in repeatedly generating temporary trait divergence. This could provide valuable insights into the mechanism of repeated, within-generation selection and its transient contribution to genetic and phenotypic variance.

## Methods

### Tag and Recapture and Environmental Data

We captured a total of 2346 *F. heteroclitus* between May 25^th^ and May 31^st^ 2018 near the Rutgers University Marine Field Station (RUMFS), NJ using wire minnow traps. Individuals were captured at four locations: a coastal basin (n = 1193) which continuously exchanges water with the open sea, and three marsh ponds (Pond 1, n = 398; Pond 2, n = 356; Pond 3, n = 399) (Fig. 1). These ponds are mostly isolated from the rest of the salt marsh except during exceptionally high tides which occur approximately 1-2x per week (Hunter et al. 2009). Every fish was weighed, measured (total length), sexed, and uniquely tagged with sequential coded wire tags (Northwest Marine Technology Inc.) injected into the dorsal musculature. After tagging, fish were released at the same location where they were captured.

Tagged fish were recaptured at each site approximately four months later between September 3^rd^ and September 7^th^ 2018. Any untagged fish were immediately released. Each pond was exhaustively fished using baited, wire minnow traps until all adult fish were captured. The coastal basin was continuously fished until less than one tagged fish was caught per 15 wire trap deployments.

In addition to the tag-and-recapture effort, we deployed HOBO® data loggers (Onset Comp.) at all locations recording temperature, conductivity, and dissolved oxygen concentration in 10-minute intervals. Data loggers were deployed between May 24^th^ and September 3^rd^ 2018 and cleaned and calibrated weekly. Raw data was processed using HOBOware Pro® and analyzed in *R*.

### Husbandry and Common Gardening

A total of 269 tagged individuals were successfully recaptured at the RUMFS and transported to our aquarium facilities at the Rosenstiel School of Marine, Atmospheric and Earth Science, Miami, FL. All fish were common-gardened in the same closed, recirculating aquarium system for 14 weeks at 20°C and 15 ppt with a 12:12 light:dark cycle. Following initial laboratory acclimation, fish were exposed to a *pseudo*-winter for 6 weeks at 10°C with an 8:16 photoperiod, then to 28°C and a 16:8 photoperiod for an additional 14-18 weeks. Hence, fish were common-gardened for a total duration of at least 8 months prior to any physiological measurements. Fish were fed *ad libidum* once per day but fasted for 24h prior to any measurement procedure. Handling, husbandry and measurement procedures were approved by the Institutional Animal Care and Use Committee (IACUC) under animal use protocol number 19-045.

### Physiological Measurements

The mass and total length of each fish were recorded before proceeding with measurement of standard metabolic rate (SMR), aquatic surface respiration (ASR) latency, critical thermal maximum (CT_max_), and substrate-specific cardiac metabolic rate (CMR).

Physiological measurements were conducted from least to most invasive in the order described above. Fish were allowed to recover for one week in between each measurement in order to minimize the effect of prior manipulation. Following physiological measurements, coded wire tags were dissected and cross-referenced with tagging data in order to identify resident (tagged and recaptured at the same site) and migrant (tagged and recaptured at different sites) individuals. All downstream statistical analyses were conducted on resident individuals only.

#### Standard metabolic rate

Standard metabolic rate was determined for each individual via automated, intermittent-flow respirometry following the high-throughput experimental setup described previously (Drown et al. 2020). Briefly, fish were acclimated to individual respiration chambers for 6 hours. Standard metabolic rate was determined by closing respiration chambers and measuring the rate of oxygen consumption for 6-minutes. Chambers were subsequently flushed and reoxygenated before starting a new measurement cycle. Multiple measurement cycles were conducted over the course of one night and oxygen consumption rates recorded for each. Individual standard metabolic rate was taken as the lower 10^th^ percentile of all measurements. Further methodological details can be found in Drown et al. (2020).

A linear mixed model was constructed modelling SMR as a function of microhabitat and the covariates mass, sex, pond ID, and their first order interactions. Day of measurement and chamber number were included as random effects. Mass was included as a linear predictor since it produced a better fit to the data than a log-scaled predictor. Data points with unusually large Cook’s distances were removed as outliers through visual inspection of the distribution of Cook’s distance. AIC-informed, stepwise regression through backward elimination was performed to determine variables with a significant effect on SMR (see Supplement 2).

#### Aquatic Surface Respiration Latency

During hypoxic conditions, diffusion at the air-water interface causes the surface layer to be more oxygenated than the water column. *F. heteroclitus* takes advantage of this by approaching the surface and ventilating its gills with surface layer water. Aquatic surface respiration (ASR) can therefore be employed to temporarily mitigate hypoxic or anoxic conditions. Here, we measure the frequency and duration of ASR in *F. heteroclitus* as a proxy for individual hypoxia tolerance.

To quantify ASR, a 40-liter experimental tank was divided into 10 equally sized compartments using plastic mesh. Individual fish were randomly assigned into compartments, and the experimental tank was screened-off in order to shield fish from any external visual stimuli. The tank was maintained fully oxygenated during preparation and held at a constant temperature of 28°C throughout the assay. An assay was started by injecting nitrogen gas into the tank at a constant flow rate. Oxygen concentration was monotonously decreased for approximately 90 minutes until a concentration of 0.7 mg/L was reached. A Vernier Inc.® oxygen probe recorded oxygen concentration throughout the assay. *F. heteroclitus* ASR behavior was recorded using two GoPro® cameras. Assay duration as well as initial and final oxygen concentration varied marginally among assays due to variance in the metabolic rates of randomly assigned individuals. Assays were therefore truncated to ensure a congruent oxygen concentration range of 6.5-0.8 mg/L during which ASR was recorded.

Video data was processed by manually time-stamping every surfacing and diving event for every fish using *MTrack* in *imageJ*. A surfacing event was recorded whenever the mouth of a fish came within less than one body depth of the surface layer. Similarly, a diving event was recorded whenever the mouth left this zone. Finally, oxygen data was merged with the time stamps and oxygen concentrations assigned to each surfacing and diving event.

Two separate statistical analyses were conducted in *R*. First, the proportional time spent performing ASR during hypoxic conditions (<2 mg/L O_2_) was calculated for each individual. A linear mixed model was constructed modelling proportional surface time as a function of microhabitat and the covariates mass, sex, pond ID, and their first order interactions. Assay ID and compartment number were included as random effects. Mass was included as a linear predictor since it produced a better fit to the data than the log-scaled predictor. Data points with unusually large Cook’s distances were removed as outliers through visual inspection of the distribution of Cook’s distance. AIC-informed, stepwise regression through backward elimination was performed to determine variables with a significant effect on proportional ASR time (see Supplement 2).

In addition to mixed linear modelling, we also performed multi-state modelling (MSM). MSM analysis is commonly used in clinical research to model the transition of patients through several disease states. Using information from multiple individuals and the timing of transitions from one state to another, the probabilities of transitioning and the average time spent in every state can be modelled, even when these are conditioned by covariates. We constructed a simple multi-state model using the *msm* package in *R* with two states: performing ASR and non-ASR. We derived the probabilities of initiating ASR and the mean time in the ASR state as a function of oxygen concentration and microhabitat. Hence, we were able to compare the readiness to engage in ASR and the time spent performing ASR between basin and pond residents. As before, mass, sex and their first order interactions were included as covariates.

#### Critical Thermal Maximum

Critical thermal maximum (*CT_max_)* was quantified by placing 10 randomly assigned fish into a 40-liter experimental tank. Starting at the acclimation temperature of 28°C, temperature was linearly increased at a rate of 0.3°C/min using an electric heating coil. An NST digital thermometer was used to monitor the temperature of the tank and record CT_max_. Critical thermal maximum was defined as the temperature at which a fish showed a loss of equilibrium and no active escape response for 5 continuous seconds. The experimental tank was maintained oxygenated and mixed throughout the assay using an air stone and a circulation pump. After reaching CT_max_, individuals were immediately removed from the tank and placed into a recovery bath. The assay was continued until all fish had reached their respective CT_max_. All assays were conducted at the same time of day. As before, a linear mixed model was constructed using the same covariates as above. Assay ID was included as a random effect. Outlier removal was performed as above and an AIC-informed, stepwise regression was conducted to determine variables with a significant effect on CT_max_ (see Supplement 2).

#### Substrate-specific cardiac metabolic rate

Substrate-specific cardiac metabolic rate (CMR) was assayed using a custom respirometer and protocol described in DeLiberto et al. (2020). Briefly, heart ventricles were dissected following euthanization by cervical dislocation. Ventricles were immediately placed into Ringer’s glucose heparin solution. Cardiac metabolic rates were assayed by measuring the oxygen consumption rates of ventricles via closed-chamber respirometry in four temperature-controlled respiration chambers. Each chamber contained a different substrate solution: 1) 5 mM glucose, 2) 1 mM palmitic acid bound to BSA (fatty acid), 3) 5 mM lactate, 5 mM hydroxybutyrate (ketones) and 0.1% ethanol (LKA) and 4) Ringer’s solution with no metabolic substrate (endogenous metabolic rate only). In addition, glycolytic enzyme inhibitors were added to all but the glucose solution in order to suppress glycolytic metabolism during the measurement of fatty acid, LKA and endogenous metabolic rates. Ventricles were rotated through each of the four respiration chambers in the order described above and oxygen consumption rates recorded for a total of 6 minutes per chamber/substrate. Cardiac metabolic rate was taken as the rate of oxygen consumption during the final 3 minutes. Four linear mixed models were constructed, one per substrate, which modelled CMR using the same fixed effects as above with ventricle mass instead of organismal mass as a covariate. Experimental batch was included as a random effect in order to account for minor differences in substrate concentration while assay time was included as a fixed effect to account for sensor drift. Outlier removal was performed as above, and an AIC-informed, stepwise regression was conducted to determine variables with a significant effect on each substrate-specific CMR (see Supplement 2).

## Author Contributions

M.A.E, M.F.O. and D.L.C. conceived the study. All authors contributed to the design of the study. M.A.E., A.N.D. and M.K.D. acquired and processed raw data. M.A.E. analyzed the data and drafted the manuscript. All authors were involved in editing and have approved submission of this manuscript. M.F.O. and D.L.C provided funding for this research.

## Ethics Statement

Fieldwork was completed within publicly available lands where no permissions were required for access. *F. heteroclitus* does not have endangered or protected status, and small marine minnows do not require collection permits for non-commercial purposes. Adult *F. heteroclitus* were captured using minnow traps under minimal stress and removed in under an hour. Tag-and-recapture and physiological assays were in compliance with and approved by the University of Miami Institutional Animal Care and Use Committee (IACUC, protocol no. 19-045).

## Acknowledgements

We would like to thank Dr. Kenneth Able, Roland Hagan, and all technicians at the Rutgers University Marine Field Station for their useful discussions regarding mummichog ecology and extensive support during fieldwork. Many thanks to Kyle Yurkow for assisting with setting up environmental sensors and tagging thousands of fish. Finally, many thanks to Liam Dorsey, Agatha Freedberg and Rebecca VanArnam for countless hours in the lab, conducting and monitoring physiological assays. M.F.O. discloses support for the research of this work from the National Science Foundation [grant number IOS 1754437] and D.L.C. discloses support for the research of this work from the National Science Foundation [grant number IOS 1556396].

## Data Availability

The environmental and physiological raw data generated and analyzed in this study are available from the corresponding author on reasonable request. Data shown in figure 5 and the corresponding source references are provided in the supplementary information files.

