## Supplemental Data for "Relentless Selection: The importance of within-generation selection in heterogeneous habitats"

**Environmental Data**

Mean environmental data for all microhabitats collected from May to September 2018.

| *Micro-habitat* | *Mean Temperature, ºC (s.d.)* | *Diurnal Temperature Variance, s.d.* | *Diurnal hours above 28ºC (s.d.)* | *Dissolved Oxygen, mg/L (s.d.)* | *Diurnal Dissolved Oxygen Variance, s.d.* | *Diurnal hours of hypoxia (s.d.)* | *Diurnal hours of anoxia (s.d.)* | *Salinity, ppt (s.d.)* | *Diurnal Salinity Variance, s.d.* |
|---|---|---|---|---|---|---|---|---|---|
| *Basin* | 22.3 (3.1) | 1.18 | 0.37 (1.27) | 5.76 (2.83) | 1.71 | 1.6 (3.1) | 0.4 (1.5) | 29.7 (1.7) | 0.78 |
| *Pond 1* | 25.7 (4.5) | 3.16 | 6.75 (4.76) | 3.33 (5.14) | 3.94 | 14.2 (5.7) | 12.2 (6.5) | 30.6 (3.4) | 1.15 |
| *Pond 2* | 26.6 (3.5) | 1.75 | 8.66 (7.50) | 3.68 (4.63) | 2.98 | 11.8 (7.1) | 8.8 (7.6) | 29.8 (2.4) | 0.48 |
| *Pond 3* | 26.3 (3.5) | 1.86 | 7.83 (7.33) | 3.40 (4.39) | 2.79 | 12.6 (6.9) | 8.9 (7.3) | 28.3 (3.0) | 0.48 |

**Linear Mixed Model Summary**

Summary statistics of linear mixed models constructed for each physiological trait. Note: microhabitat is not a statistically significant fixed effect in cardiac metabolic rate with fatty acid substrate but is being shown here for its marginal significance.

| *Trait Measured* | *Sample size (before outlier exclusion)* | *Random Effects* | *Significant Fixed Effects* | *Effect Size* | *p-value* |
|---:|---:|---:|---:|---:|---:|
| *Standard Metabolic Rate* | 181 (188) | *respiration_chamber*<br>*assay_ID* | *microhabitat (Pond)*<br>*mass (g)* | -0.33<br>0.34 | 0.024<br><< 0.001 |
| *Aquatic Surface Respiration (prop. time at surface)* | 110 (125) | *ASR_chamber*<br>*assay_ID* | *microhabitat (Pond)*<br>*mass (g)*<br>*sex (male)* | -0.07<br>-0.01<br>-0.12 | 0.013<br>0.023<br>0.001 |
| *Critical Thermal Maximum* | 160 (164) | *assay_ID* | - | - | - |
| *Cardiac Metabolic Rate (Glucose)* | 141 (147) | *assay_ID* | *microhabitat (Pond)*<br>*ventricle_mass (µg)*<br>*assay_time* | -3.74<br>1.42<br>-0.46 | 0.004<br><< 0.001<br>< 0.001 |
| *Cardiac Metabolic Rate (FA)* | 144 (146) | *assay_ID* | *microhabitat (Pond)*<br>*ventricle_mass (µg)*<br>*assay_time* | *-1.77*<br>1.16<br>-0.48 | *0.053*<br><< 0.001<br><< 0.001 |
| *Cardiac Metabolic Rate (LKA)* | 137 (144) | *assay_ID* | *ventricle_mass (µg)*<br>*assay_time* | 1.34<br>-0.55 | << 0.001<br><< 0.001 |
| *Cardiac Metabolic Rate (Endogenous)* | 133 (140) | *assay_ID* | *ventricle_mass (µg)*<br>*assay_time*<br>*ventricle_mass*sex* | 0.49<br>123.55<br>15.97 | << 0.001<br><< 0.001<br><< 0.001 |

**Trait Correlations**

Matrix of pairwise correlations between all physiological traits. The lower half holds scatter plots of the residuals, the upper half shows the Pearson correlation coefficient between residuals and the diagonal shows the density distribution of residuals for each trait. Significant Pearson correlation coefficients are indicated with asterisks.

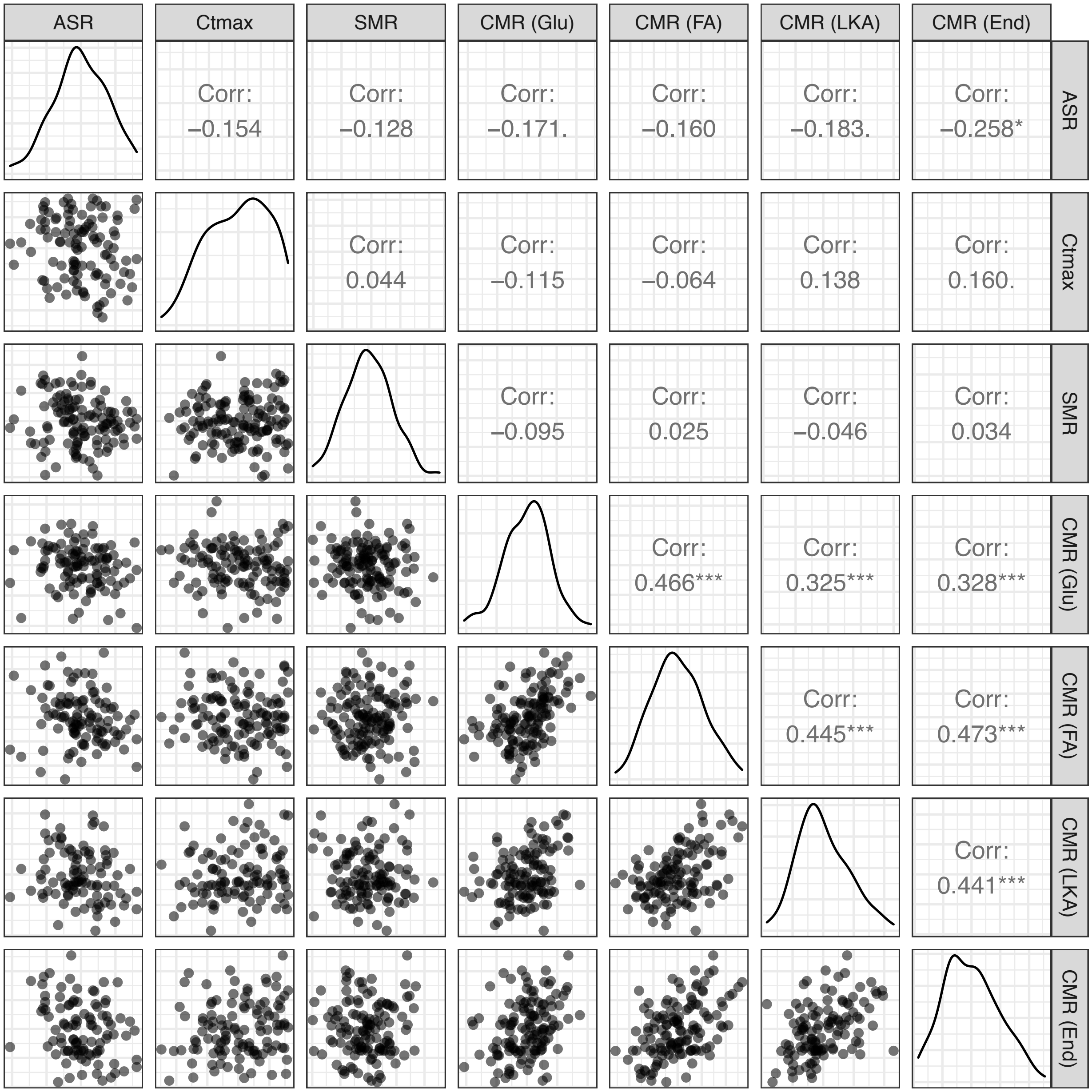

**Partial Trait Correlations**

Matrix of partial correlation coefficients for each pairwise comparison for all physiological traits. Partial correlations show the unbiased correlation between two traits by removing the effect of all other traits.

| | ***ASR*** | ***$CT_{max}$*** | ***SMR*** | ***CMR (Glucose)*** | ***CMR (FA)*** | ***CMR (LKA)*** | ***CMR (Endo.)*** |
|---|---|---|---|---|---|---|---|
| ***ASR*** | 1.000 | -0.048 | -0.037 | -0.002 | -0.014 | 0 | -0.078 |
| ***$CT_{max}$*** | -0.021 | 1.000 | 0.001 | -0.329 | -0.037 | 0.025 | 0 |
| ***SMR*** | -0.005 | 0.006 | 1.000 | -0.008 | -0.01 | 0.19 | 0.010 |
| ***CMR (Glucose)*** | -0.001 | -0.095 | -0.221 | 1.000 | 0.217 | -0.005 | 0.018 |
| ***CMR (FA)*** | -0.042 | -0.066 | -0.010 | 0.184 | 1.000 | 0.055 | 0.030 |
| ***CMR (LKA)*** | -0.001 | 0.025 | 0.001 | -0.002 | 0.061 | 1.000 | 0.069 |
| ***CMR (Endo.)*** | -0.079 | 0 | 0.013 | 0.016 | 0.204 | 0.107 | 1.000 |

**Metanalysis Publications List**

Data used to construct the distribution of standardized trait divergence (Figure 5) was taken from the following list of publications. Data was only included if a significant trait difference between two populations was detected in a common-garden or reciprocal transplant design. Note that the majority of data was taken directly from two major meta-analyses by Grainger & Levine (2022) and Leimu & Fischer (2008)(see below).
